# Development of visible light-sensitive human OPN5 via single amino acid substitution

**DOI:** 10.1101/2025.05.19.654910

**Authors:** Yusuke Sakai, Richard J. McDowell, Robert J. Lucas

## Abstract

Animal opsins, light-sensitive G-protein-coupled receptors (GPCRs), serve as primary light receptors in animals, supporting vision and providing light-dependent regulation of physiology and behaviour. Animal opsins also show great potential to be used as optogenetic tools that regulate cellular G protein signalling with light. Opn5 is a UV-sensitive “non-visual” opsin that is expressed in a wide range of tissues throughout body. Although Opn5 could be a good template for the development of optogenetic tools applicable to tissues outside of the eye because of its broad expression, its sensitivity to poorly tissue-penetrating UV light poses challenges for its application. In this study, we focused on human OPN5 (hOPN5) to attempt to identify amino acid(s) responsible for the UV sensitivity and to shift the spectral sensitivity to visible light. Sequence alignment across UV-sensitive Opn5s identified a conserved lysine reside (Lys91) at a position implicated in spectral tuning in invertebrate opsins. We applied site-directed mutagenesis to replace this residue with neutral (alanine) or acidic (glutamate or aspartate) amino acids. Heterologous action spectroscopy of these mutants revealed substantial shifts in spectral sensitivity (55-63nm) toward visible wavelengths. Our findings identify Lys91 as a key spectral tuning site in hOPN5 and provide visible-light sensitive versions of this protein as a candidate for optogenetic applications.

## Introduction

Animal opsins are light-sensitive G protein coupled receptors that convert light into intracellular G protein signalling. In vertebrates, opsins extend beyond the well-studied rod and cone photopigments underlying vision to encompass various “non-visual” opsins identified from tissues outside photoreceptor cells. Opn5 (neuropsin) is a type of the non-visual opsin first identified in the genomes of human and mouse [1]. Mouse Opn5 is expressed in a wide range of tissues including brain and skin [1,2] and has been reported to mediate various light-dependent physiologies such as direct photoentrainment of local circadian clocks in cornea [3] and skin [4] and the violet-light dependent suppression of brown adipose tissue (BAT) activity [5]. In non-mammalian vertebrates, quail Opn5 is expressed in the hypothalamus paraventricular organ (PVO) where it contributes to regulation of seasonal cycle of reproduction [6]. In the Japanese rice fish (medaka), Opn5 expressed in pituitary melanotrophs regulates the short-wavelength-light dependent release of melanocyte-stimulating hormone (MSH) and subsequent pigmentation in the skin [7].

In addition to such physiological importance, the Opn5 family has also attracted attention for its potential for generating optogenetic tools that regulate intracellular G protein signalling with light. A recent study has shown that human OPN5 (hOPN5) selectively activates the Gq-type G protein without promiscuous activation of other G proteins such as Gi [8] which most other Gq-coupled opsins including melanopsins [9,10] and jumping spider Rh1 [11,12] activate. hOPN5 as a Gq-selective optogenetic tool has been applied *in vivo* to murine cardiomyocytes to modulate heart beat rate and smooth muscle cells in various organs such as the small intestine and stomach in mice to induce muscle contraction with light via Gq signalling pathways [8,13].

Although hOPN5 is becoming increasingly valuable in the field of optogenetics, several challenges remain to be addressed for its widespread application. One of the most important problems is that it is primarily sensitivity to UV light, which has much lower penetration of biological tissues than visible light due to higher absorbance and scattering. Homologues of hOPN5 from across the vertebrates (the so called ‘Opn5m’ sub-family; supplementary material, figure S1) form UV-absorbing pigments with λ_max_ at 360 ∼ 380 nm upon binding 11-*cis* retinal in the dark [2,14]. The highly conserved UV sensitivity among these Opn5m opsins suggests the existence of shared amino acid residue(s) responsible for the UV sensitivity. In this study, we set out to identify such responsible amino acid residue(s) for the UV sensitivity of hOPN5. We applied sequence alignment and homology modelling to identify conserved residues close to the chromophore binding pocket with potential to provide UV sensitivity. We then applied site directed mutagenesis and heterologous action spectroscopy [9,15] in HEK293 cells to determine their involvement in this property. We find that a lysine residue, Lys91 (bovine rhodopsin amino acid numbering), in helix II is necessary to make hOPN5 UV-sensitive. Based on this result, we discuss the potential use of hOPN5 K91 mutants as visible light-sensitive Gq-manipulating optogenetic tool.

## Results

We first measured light-dependent Ca^2+^ increases in wild type hOPN5-transfected HEK293T cells via an aequorin-based bioluminescent assay (figure 1a) and confirmed that it induced light intensity-dependent Ca^2+^ increases in the cells (figure 1b). The irradiance-response curves for six spectrally distinct stimuli showed that the wild type hOPN5 had a higher sensitivity to stimuli rich in UV than visible light (figure 1c). This finding confirms that hOPN5 forms UV-sensitive pigments in the HEK293T cell environment to drive Gq-mediated Ca^2+^ responses upon light absorption, as previously reported [2,8,14].

**figure. 1.**
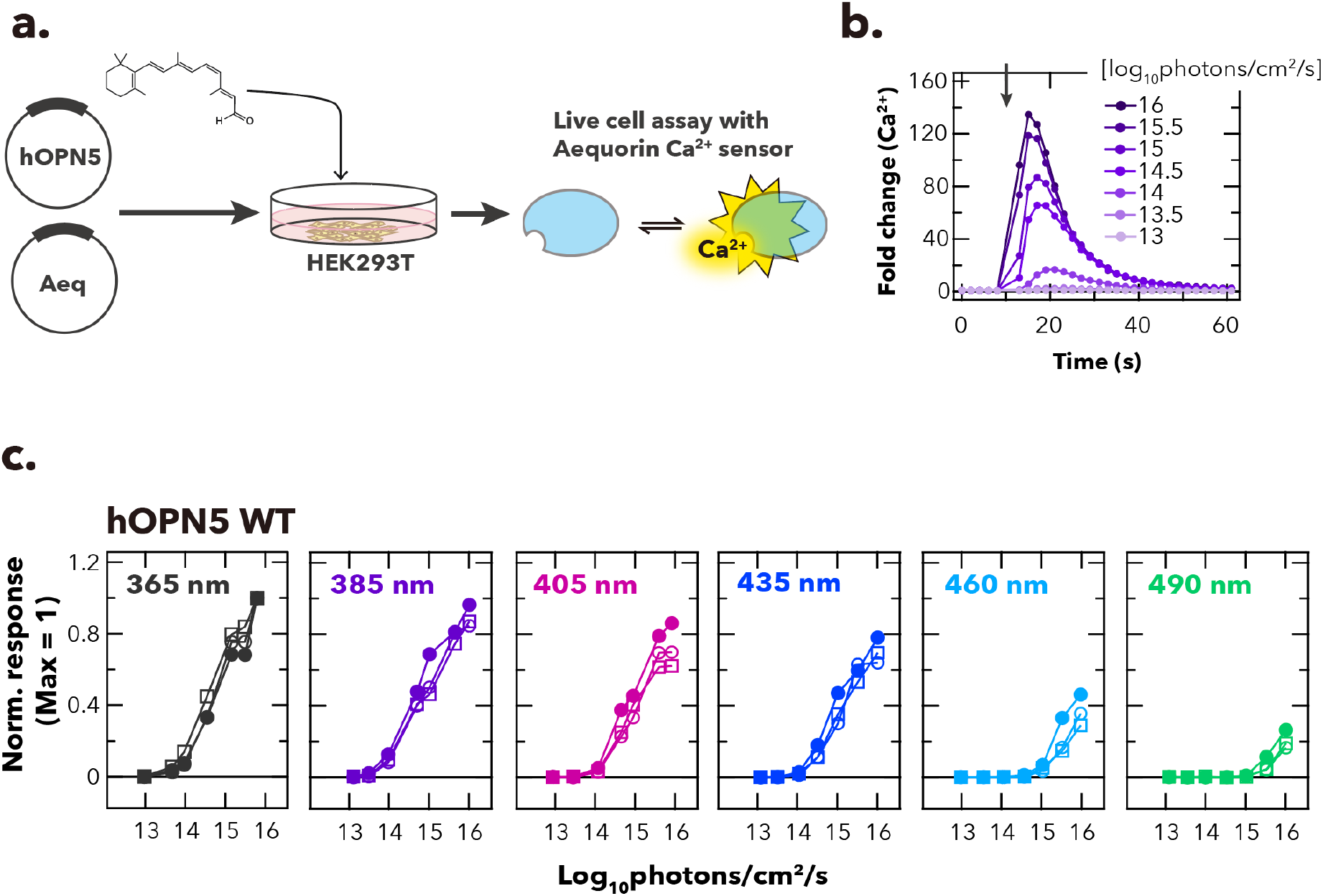
Measurement of light-induced Ca^2+^ response in HEK293T cells transfected with wild type hOPN5. **a**. Schematic of bioluminescence-based Ca^2+^ assay using aequorin Ca^2+^ biosensor. HEK293T cells were transfected with hOPN5- and aequorin-expression vectors and incubated with 11-*cis* retinal. Bioluminescence from aequorin Ca^2+^ sensor was recorded using a plate reader. **b**. Time course of light-evoked Ca^2+^ increase in the hOPN5-expressing cells. Luminescence values were normalised to the baseline (= 1). The cells were stimulated with various intensities of 385-nm light (1 s duration) at the time indicated by the black arrow in the graph. **c**. Irradiance-response curves of wild-type hOPN5-expressing cells for six spectrally distinct light stimuli (numerical values at top left depict wavelength at peak photon flux). The peak Ca^2+^ responses achieved for each stimulus were normalised to the maximum value observed across all wavelength & intensity combinations (as 1) for each independent experiment. Each replicate is indicated by different shapes of symbols (n = 3).

To determine the amino acid residue(s) responsible for the UV sensitivity of hOPN5, we searched animo acids which are conserved across the Opn5m sub-family and located near the Schiff base based upon AlphaFold3 structural modelling [16]. The most promising candidate was a conserved Lys (position 91) in helix II of Opn5s (figures 2a, b). Replacement of Lys residues at an equivalent location in UV-sensitive *Drosophila melanogaster* Rh3 and Rh7 opsins and *Platynereis dumerilii* c-Opsin1 have successfully produced shifts to visible wavelength sensitivity [17–19].

**figure. 2.**
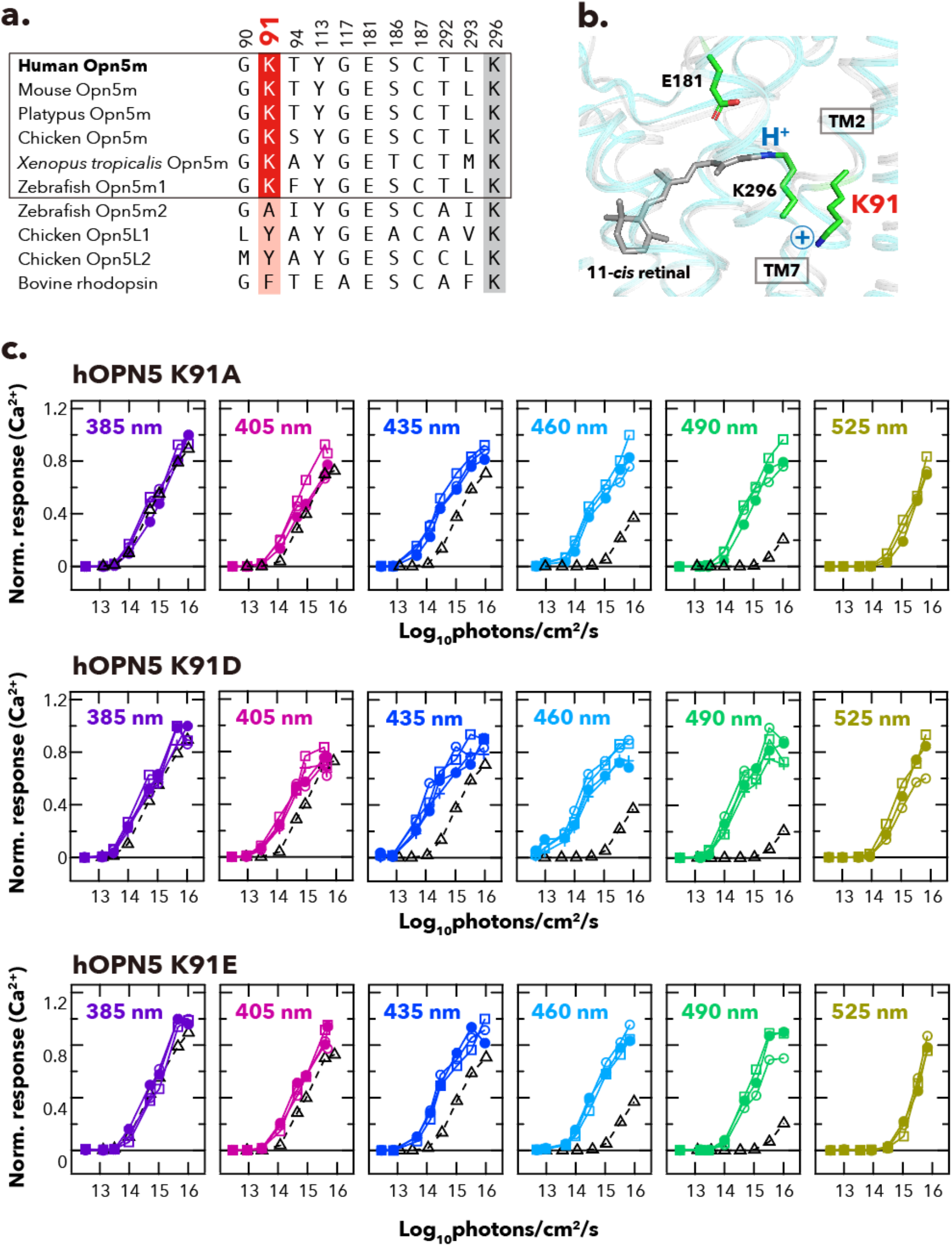
Lys91 mutants hOPN5 show enhanced sensitivity to visible light. **a**. Comparison of amino acid residues near the retinylidene Schiff base among opsins in Opn5 group and bovine rhodopsin. **b**. The Schiff base environment in hOPN5 depicted according to Alpha-fold3 3D model of hOPN5 (cyan) and X-ray crystal structure of bovine rhodopsin (PDBID: 1U19, grey). **c**. Irradiance-response curves of hOPN5 K91A, K91D and K91E are shown for six spectrally distinct light stimuli (numerical values at top left depict wavelength at peak photon flux). The peak Ca^2+^ responses achieved for each stimulus were normalised to the maximum value observed across all wavelength & intensity combinations (as 1) for each independent experiment. Each replicate is indicated by different shapes of symbols (n = 3). Black open triangles with dashed lines indicate the mean values of wild type hOPN5 (reproduced from figure 1c).

To determine the contribution of Lys91 to UV sensitivity of hOPN5, we substituted the residue with amino acids having neutral (alanine) or negatively charged (glutamic/aspartic acids) side chains and examined their spectral sensitivity of the resultant K91 mutants. We confirmed that the K91 mutants increased intracellular Ca^2+^ light-dependently as observed in the wild type opsin. Construction of irradiance-response curves for spectrally distinct stimuli showed that the K91 mutants had higher sensitivity to visible light compared to the wild type (figure 2c). The λ_max_ values of wild type and K91A mutants of hOPN5 were estimated by applying a nonlinear optimisation algorithm with bootstrapping to the experimentally obtained light intensity-Ca^2+^ response dataset [15] (see supplementary material, figure S2 and Material and Methods section for details). The estimated λ_max_ values were 389 nm for wild type (similar to that reported elsewhere), while K91A, K91D, and K91E mutants of hOPN5 had λ_max_ around 447 nm, 452 nm, and 444 nm respectively (figure 3). These results indicate that a single amino acid mutation at Lys91 is sufficient to make hOPN5 visible-light sensitive.

**figure. 3.**
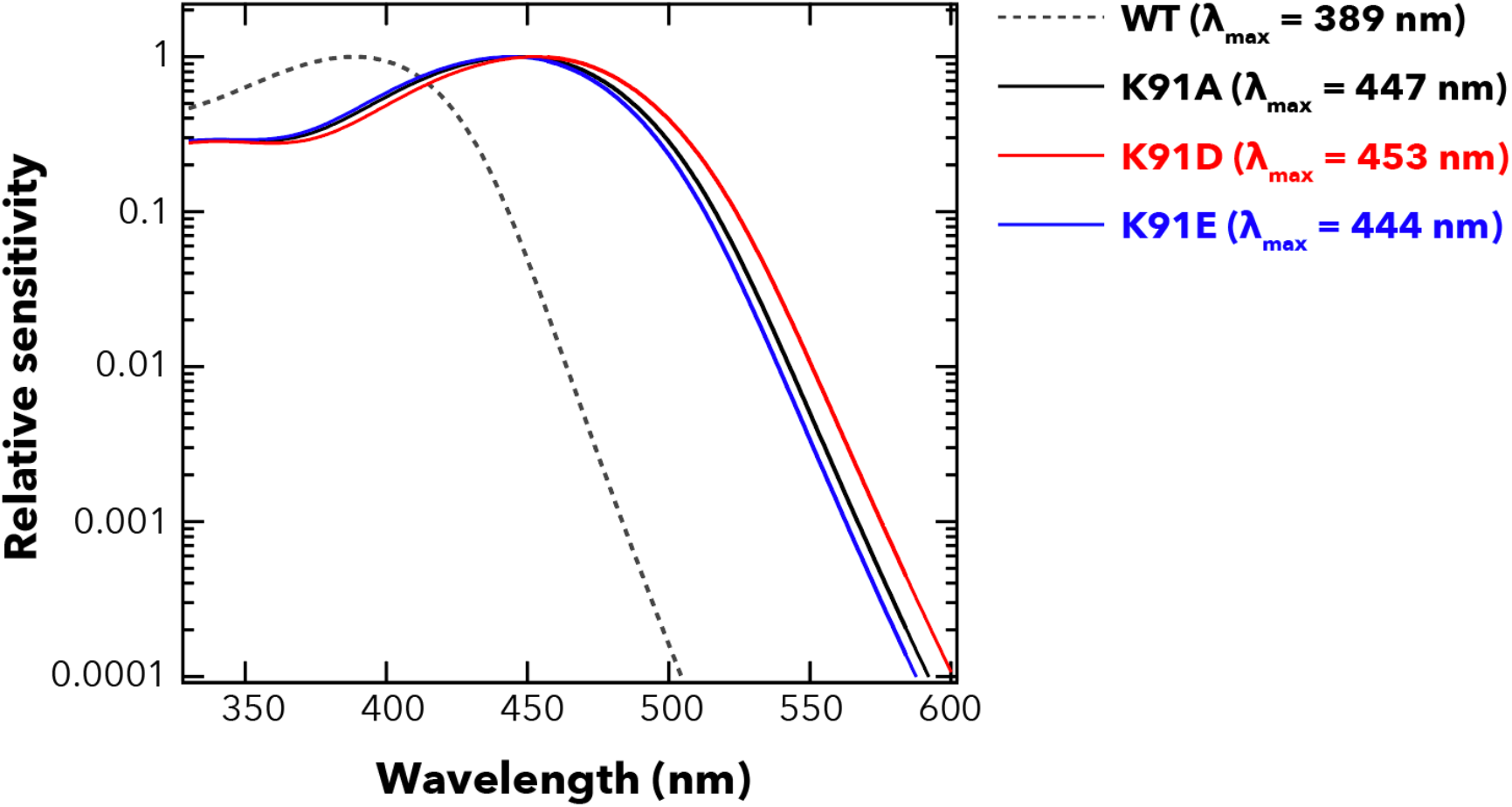
Predicted spectral sensitivity functions for wild type and Lys91 mutants of hOPN5. Curves show Govardoskii opsin pigment templates that best predict the set of irradiance-response curves depicted in figures 1 and 2 for WT (black dotted), K91A (black), K91D (red) and K91E (blue) mutants.

## Discussion

The visible light shift we observed in the K91A, K91D & K91E mutants implicates Lys91 as a critical residue in defining UV sensitivity for human OPN5. Opsins can absorb visible light only when the chromophore retinal is covalently bound to the opsin via a protonated Schiff base. Consequently, the most parsimonious explanation for the role of Lys91 in UV sensitivity is that the positive charged side chain of this residue destabilises the Schiff base protonation. Such a mechanism has been suggested to explain the contribution of Lys residues at sites 90 and 94 in *D. melanogaster* Rh3 and Rh7 and *P. dumerilii* c-Opsin1, respectively, to UV sensitivity of these opsins [17–19]. Moreover, in teleost parapinopsins (non-visual opsins phylogenetically close to vertebrate rod and cone opsins), introduction of helix II from UV-sensitive PP1 is sufficient to provide a λ_max_ at UV light region to the visible light (blue)-sensitive PP2 [20] and also, a report has demonstrated that G90K mutation in bovine rhodopsin shifts the λ_max_ to UV region (384 nm) [21]. Thus, across diverse opsins, residues in helix II appear to define UV sensitivity. However, according to the Alpha-fold3 3D model of hOPN5 aligned with the crystal structure of bovine rhodopsin (figure 2b), the side chain of Lys91 in hOPN5 is about 8Å away from the Schiff base. It remains to be determined how such a distant residue could destabilise Schiff base protonation. Furthermore, sequence alignments across more distantly related Opn5s (supplementary material, figure S1) reveals UV sensitive opsins (Opn5m2 and Opn5L2)[22,23] that lack Lys91 (figure 1a), indicating that these Opn5 homologs have different spectral tuning mechanisms from Opn5m for their UV sensitivity.

The possible immune response against optogenetic protein itself has been reported as well as against AAV-mediated gene transfer [24]. Mammalian non-visual opsins including Opn3, Opn4, and Opn5 [25] that exhibit broad tissue expression represent an attractive strategy to reduce the likelihood of such immune responses; therefore, they have a significant advantage for both experimental and therapeutic optogenetic applications. hOPN5 has generated particular interest due to its specific activation of Gq-type G proteins [8,26] and successful *in vivo* applications of hOPN5 have been demonstrated, such as light-dependent control of muscle contraction across a wide range of muscle tissues [8,13]. The visual light-sensitive version of hOPN5 in this study have the potential to be applied to deep tissues where UV light penetration is limited and thus could be a good template upon which to develop general Gq-manipulating optogenetic tools. As there has been accumulating knowledge about spectral sensitivity tuning within the visible-light range [27–29], engineering of further red-shifted mutants of hOPN5 could prove fruitful in future studies.

## Material and Methods

### Construction of opsin expression vectors

mRNA sequence of hOPN5 was obtained from NCBI GenBank (NM181744). We designed sequences containing the open reading frames of wild type and mutants of hOPN5 tagged with rho 1D4 epitope sequences (ETSQVAPA) at the C-terminus. These DNA sequences were then synthesised using gene synthesis service of Twist Bioscience. The synthesised DNA fragments were inserted into the HindIII/NotI-digested pcDNA3.1 mammalian expression vector (Thermo Fisher Scientific) using NEB HiFi Assembly (New England Biolabs).

### Bioluminescence-based Ca^2+^ measurements in HEK293T cells

HEK293T cells were cultured in Dulbecco’s modified Eagles medium – high glucose (Sigma-Aldrich) with 10% foetal bovine serum (FBS) and 1% Penicillin/Streptomycin. Cells were seeded on 12-well plates (∼500,000 cells per well) and transiently transfected ∼24 h later with expression vectors for each target opsin (500 ng) and genetically encoded bioluminescent Ca^2+^ reporter, mtAequorin (500 ng) using Lipofectamine 2000 Transfection Reagent (Thermo Fisher Scientific) as described previously [9,15]. Four to six hours after transfection, 10 µM 11-*cis* retinal (National Eye Institute, National Institutes of Health) was added to the cells. The cells were subsequently added to white clear-bottom 96-well plates for incubation at 37°C overnight. The following day, the cells were incubated in Leibovitz’s L-15 Medium (Thermo Fisher Scientific) containing 10 µM Coelenterazine-h (Promega), 10 µM 11-*cis* retinal, and 1% FBS in the dark at room temperature for 2 h before recording luminescence. For each recording, baseline luminescence was recorded for 10 s and then the cells were stimulated with 1-s light flash of varying intensities (12.5 to 16 log_10_photons/cm^2^/s) at one of distinct wavelengths (365 nm, 385 nm, 405 nm, 435 nm, 460 nm, 490 nm, and 525 nm). Luminescence signals were measured every 2 s using a plate reader (Optima FLUOStar, BMG Labtech) and an external light source (CoolLED PE-4000, CoolLED) via liquid light-guide with a set of neutral density filters (Thorlabs) was used to illuminate the cells.

### Calculation of wavelength of maximum sensitivity for each opsin

Based on irradiance-response curves obtained from the aequorin Ca^2+^ assay, we determined the λ_max_ values for each opsin using a nonlinear curve fitting with bootstrapping approach as described previously [15]. Briefly, we first calibrated the measured irradiance (log_10_photons/cm^2^/s) according to the Govardovskii opsin pigment template [30] with a given λ_max_ value (namely, each measured irradiance response curve was multiplied by a template absorption spectrum having a given λ_max_ value) and obtained the effective light intensity. Then, a 5-parameter logistic model was fitted with the normalised Ca^2+^ increase as the dependent variable and the effective light intensity as the indepedent variable using the *drm* function from the *drc* package in R. The residual sum of squares (RSS) extracted from each fitting were passed to the optimisation algorithm (*optim* function in R using the Brent search method) to identify the λ_max_ value (within a 350 – 550 nm range) that would minimise the RSS. Bootstrapping (100 iterations) was applied, in which the optimisation procedure was performed for each dataset constructed based on resampling with replacement from the original dataset, and the average value of the 100 replicates were used as the λ_max_ estimate. For λ_max_ estimation, we used irradiance-response datasets at 365 nm, 385 nm, 405 nm, and 435 nm for wild type hOPN5 and 385 nm, 405 nm, 435 nm, 460 nm, and 490 nm for K91 mutants hOPN5, where Ca^2+^ responses were nearly saturated. All analyses were performed in R version 4.4.3 [31].

### Molecular phylogenetic tree inference

Amino acid sequences of opsins obtained from NCBI GenBank database were aligned using MAFFT [32] and trimmed by TrimAl [33] with “gappyout” function. The maximum likelihood tree was reconstructed using RAxML-NG v.1.2.0 [34] assuming the LG + I + G4 + FC model, which was selected using ModelTest-NG v.0.2.0 [35]. The ML branch supports were obtained by 500 bootstrap resamplings.

## Acknowledgement

We thank National Eye Institute, National Institutes of Health for providing 11-*cis* retinal.

## Author contributions

YS: conceptualisation, investigation, formal analysis, data curation, visualisation, writing—original draft, writing—review and editing.

RJM: methodology, investigation, writing—review and editing

RJL: conceptualisation, writing—original draft, writing—review and editing, supervision, project administration, funding acquisition.

## Supplementary materials

**figure S1.**
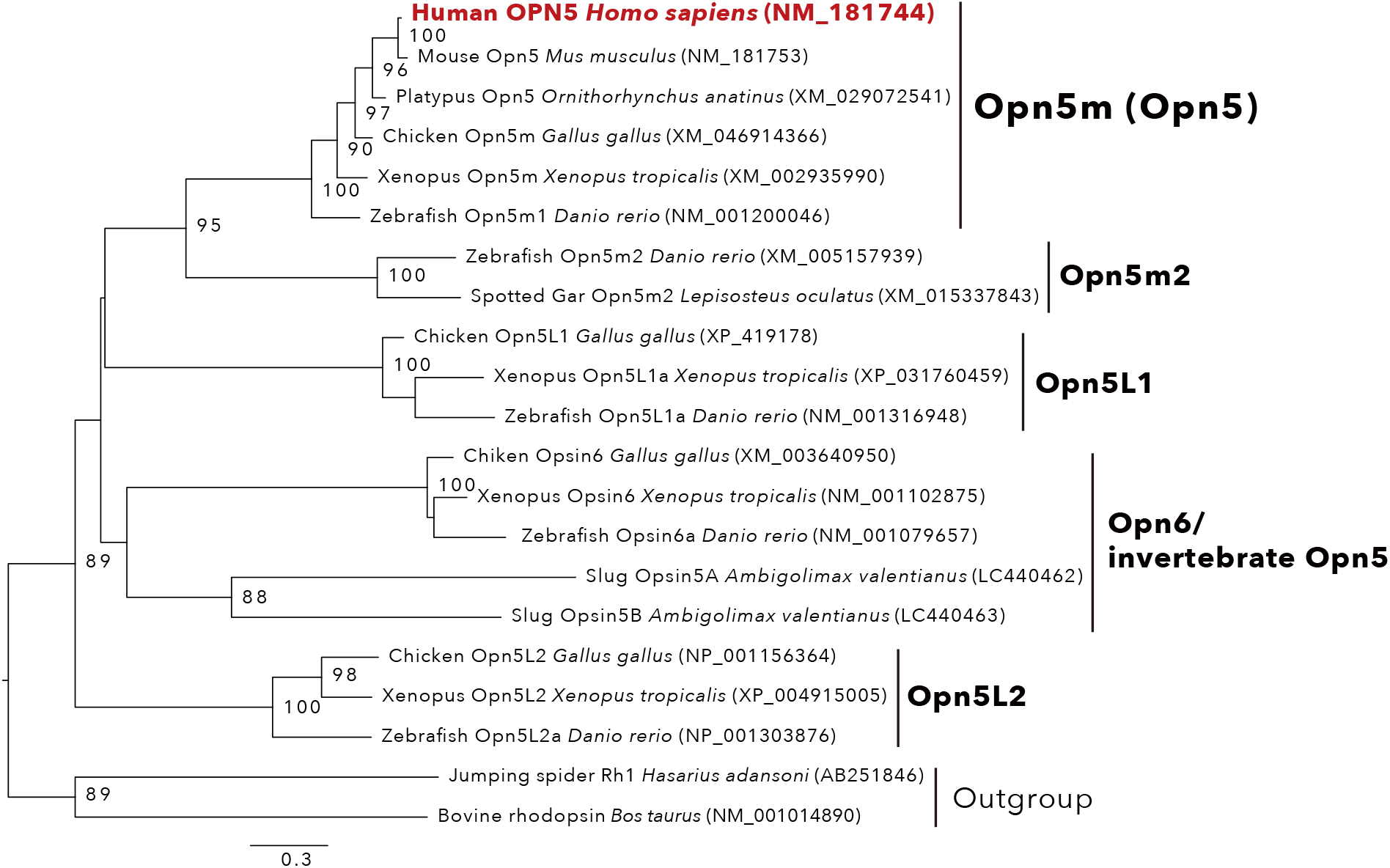
Maximum likelihood tree of Opn5 group. Numbers at the nodes indicate bootstrap support values (≥ 80% are indicated). A scale bar = 0.6 substitutions per site.

**figure S2.**
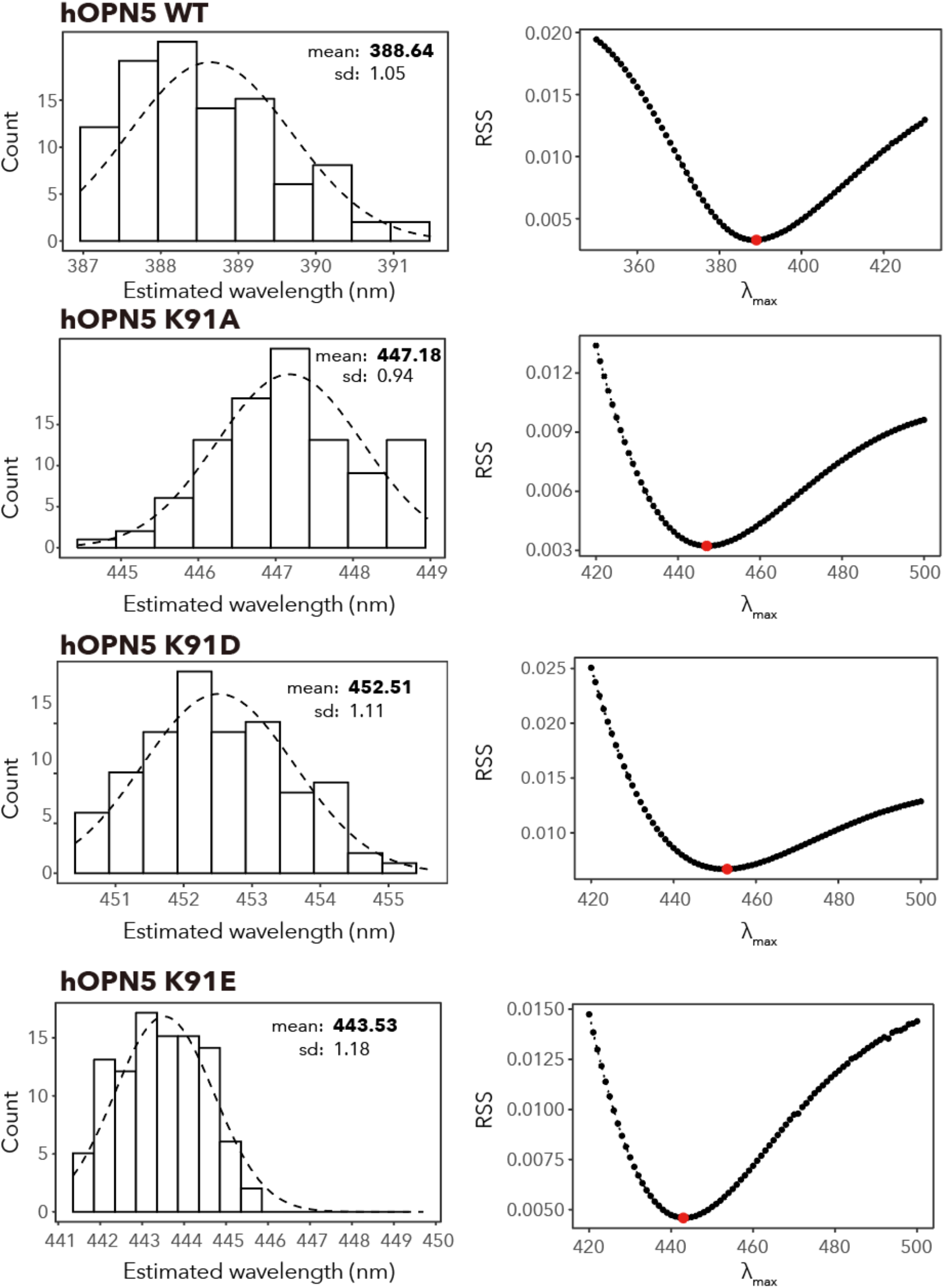
Estimation of spectral sensitivities for wild type and Lys91 mutants of hOPN5 by a nonlinear optimisation with bootstrap samplings. (Left panels) Histograms of λ_max_ values calculated from 100 bootstrap samples. (Right panels) Residual sum of squares (RSS) values showing the goodness-of-fit of the 5-parameter logistic curve fitting to the cell responses (response variable) and the effective light intensity (explanatory variable) calculated under λ_max_ values (wavelength ranges: 350 nm – 430 nm for wild type and 420 nm – 500 nm for Lys91 mutants). The RSS value calculated under the predicted λ_max_ for each pigment were shown in red circle.

